# Zoonotic Tick Borne Pathogens in Ticks at Human-Wildlife-Livestock interfaces in Kwale County, Kenya

**DOI:** 10.1101/2022.03.07.483224

**Authors:** Salim Kobo Godani, Nelson Menza C., Margaret Muturi W.

## Abstract

**Background:** Ticks are vectors that can harbor pathogens like viruses, protozoa and bacteria that can cause zoonotic diseases in human. Human gets infected through tick bites where the pathogens are passed to the human blood. One tick bite can lead to transmission of more than one disease. Ticks are obligate hematophagous, infest most vertebrates globally. Lack of surveillance information about the tick-borne pathogens and diseases has made it impossible to assess its impact to the community.

**Materials and Methods:** We morphologically identified ticks collected from two game reserves in Kenya, locations near the Shimba Hills National Reserve (SHNR) and Tsavo National Reserve (TNR). We identified tick-borne pathogens by HRM analysis and sequencing of specific RT-PCR products of Anaplasma, Ehrlichia, and Rickettsia.

**Results:** A total of 317 (281 adult ticks and 36 nymphs) was collected at near Tsavo National Reserve in Taita Taveta County, which includes seven species. *Amblyomma* was the most sampled genus with *Amblyomma gemma* being the most sampled species (n=135, 42.6%). Other *Amblyomma* species sampled was *Amblyomma Variegatum* (n= 40, 12.62%). Greatest species diversity was identified in *Rhipicephalus* genus with four species identified that includes; *Rhipicephalus appendiculatus* (n=44, 13.9%), *Rhipicephalus Averts* (n=1, 0.31%), *Rhipicephalus Decoloratus* (n=5, 1.6%), *Rhipicephalus Pulchellus* (n= 91, 28.7%). A single species of *Hyalomma* sp. was sampled. From near Shimba Hill game reserve (SHNR), a total of 240 adult’s ticks were sampled that comprises of eight species. *Amblyomma* was the most sampled genus and again *Amblyomma gemma* being the most sampled species (n=156, 65 %). Other *Amblyomma* species sampled includes; *Amblyomma Lepidum* (n= 5, 2.1 %), *Amblyomma Variegatum* (n= 15, 6.3 %). Greatest species diversity was also identified in *Rhipicephalus* genus with four species identified that includes; *Rhipicephalus appendiculatus* (n=18, 7.5 %), *Rhipicephalus Averts* (n=6, 2.5 %), *Rhipicephalus Decoloratus* (n=4, 1.7 %), *Rhipicephalus Pulchellus* (n= 34, 14.2 %). The least sampled species was a single species of *Hyalomma* Scupense (n=2, 0.83 %). At near Tsavo National Reserve (TNR) in Taita Taveta County, a total of three pools of *Rhipicephaline appendiculatus* were positive for *Theileria parva*, two pools of *Rhipicephaline evertsi* for *Anaplasma platys* and one pool of *Amblyomma variegatum* nymphs for *Rickettsia africae*. From near Shimba Hill game reserve (SHNR), Kwale County, *Rickettsia africae* pathogen was detected in two pools of *Am. variegatum* and one pool of *Am. Gemma. Rickettsia sp*. and *Anaplasma sp*. were detected in *Am. Gemma* and *Rh. evertsi* respective. *R. aeschlimannii* was detected in a pool of *Am. Gemma*. These findings highlight the risk of transmission of zoonotic *R. africae* and unclassified *Rickettsia* spp. to humans causing African tick-bite fever and other spotted-fever group rickettsioses.

## Introduction

Arthropod vectors like ticks can harbor viruses, bacteria, and protozoa which can cause diseases in wildlife and livestock (1, 2). They are obligate hematophagous that parasitize most vertebrates in almost every region in the world (3). Ticks are second to mosquitoes as vectors with 800 species which depends on the blood of most vertebrates and cause disease in human (3). Ticks belong to class *Arachnida*, only two families, the *Ixodidae* (hard tick) and *Argasidae* (soft ticks) have medical importance. These families can be differentiated by the presence of the scutum, hard ticks possess the scutum and soft ticks lack the scutum (4). Humans get the infection through a tick bite, direct contact from viremia tissues of humans or animals, and the disease is characterized by hemorrhagic (5) and a single tick bite can transmit more than one pathogen (6).

The vector-borne infections have been a problem for both animals and humans causing black death plagues in Europe and yellow fever in America in 1th centaury (7). Tick-borne diseases include: *Ehrlichiachaffeensis, Ehrlichiaewingii, E. canis, E. ruminantium*, and *Anaplasma phagocytophilum*, and transmission is by ticks (8). Tick-borne diseases are amongst neglected tropical diseases leading to high prevalence and distribution morbidity (9). Lack of the tick-borne zoonotic information in sub-Saharan Africa made it impossible to assess their impact (9). Hard ticks harbor intracellular bacteria that show mutualistic and pathogenic associations to the host, the bacteria can be transmitted to mammals including humans (10). The most important tick-borne disease in Kenya includes Theileriosis, anaplasmosis, babesiosis, and heartwater (11). A study was done along with Lake Victoria and Lake Baringo about the ticks molecular and pathogenic diversity by Omondi *et al* (12) reported pathogens like canine ehrlichiosis (E. ruminantium, Ehrlichia canis, and Ehrlichia sp). anaplasmosis (Anaplasma ovis, Anaplasm a platys and A. bovis) rickettsiosis (Rickettsia aeschlimannii) in ticks.

There is limited knowledge of circulating tick-borne pathogens of medical importance in their biological vectors at human-wildlife-livestock interface in Kenya due to limited resources. This study aimed at addressing the gap in knowledge on the zoonotic tick-borne pathogens circulating at human-wildlife-livestock interfaces in the Tsavo National Reserve (TNR) and the Shimba Hills National Reserve (SHNR), Kenya, by surveying ticks’ diversity and determining tick-borne pathogens.

## Materials and Methods

### Study Area

Sampling approval was obtained from the Kenya’s Directorate of Veterinary Services and Ministry of Health. Informed consent was obtained from the livestock’s owner before sampling of the ticks at near Tsavo and Shimba Hills National Reserves, Kenya. The two selected human-wildlife-livestock interfaces were selected based on encroachment human settlement by the pastoral community and reported cases of tick-borne pathogens of economic and public health burden. The Tsavo national park is a protected area in Kenya, which got its name from the Tsavo River. The park borders Chyulu hills National Park and Mkomazi game reserve of Tanzania. The park is divided by the highway road that runs from Nairobi to Mombasa into Tsavo East national park (13,747 Km^2^) and Tsavo West national park (9,065 Km^2^).

The two National Parks have different ecosystems that ranges from the flat undulating open savannahs on the East to the hilly volcanic mountains on the West. Tsavo East national park is a home to a variety of wildlife species, including red dusty Elephants *(Loxodonta)*, lions *(Panthera leo)*, waterbucks *(Kobus ellipsiprymnus)*, kudu *(Tragelaphus strepsiceros)*, crocodiles *(Crocodylinae)*, Hippopotamus *(Hippopotamus amphibious)*, leopards *(Panthera pardus)*, zebras *(Equus quagga)*, giraffes *(Giraffa)*, and gerenuk *(Litocranius walleri)*. It also hosts over 500 bird species that are recoded at the park. The SHNR is a wildlife protected area in Kwale County. It is the largest forest area in east African and home to a variety of wildlife species, including endangered sable antelopes *(Hippotragus niger)*, Elephant *(Loxodonta)*, giraffes *(Giraffa)*, leopards *(Panthera pardus)*, waterbuck *(Kobus ellipsiprymnus)*, bush duiker *(Sylvicapra grimmia)*, buffalo *(Syncerus caffer)*, bush pigs *(Potamochoerus larvatus)*, African bushbabys *(Galago moholi)*, coastal black-handed titi *(Callicebus melanochir)*, black and white colobus *(Colobus guereza)*, Sykes’ monkeys *(Cercopithecus albogularis)*, greater galago black-faced vervet monkeys *(Chlorocebus pygerythrus)*, serval cats *(Leptailurus serval)*, warthogs *(Phacochoerus africanus)*, and bushbucks *(Tragelaphus scriptus)*.

### Tick collection and identification

Sampling approval was obtained from the Kenya’s Directorate of Veterinary Services and Ministry of Health. Informed consent was obtained from the livestock’s owner before sampling of the ticks. Ticks were collected from restrained cattle hide using sterile forceps and placed into labeled falcon tubes. The tubes were then plugged with cotton swabs and transported in liquid nitrogen to the testing lab. They were stored at −80°C until when analysed (12). The ticks were identified to the genus and/or species level as per the morphological keys (13) using sterile forceps and sterile petri-dish while on sterile gloves. Ticks were pooled according to species, sex and sampling sites into groups of 1–11 for adults and 1–20 for nymphs as described by Oundo *et al*. (14).

### Laboratory Procedure

#### DNA extraction

Ticks were homogenized in a 1.5-ml sterile microcentrifuge tubes containing 750 mg of 2-mm yttria-stabilized zirconium using a handheld battery operated homogenizer oxide beads (Glen Mills, Clifton, NJ). DNA was extracted from the homogenate using the DNeasy Blood and Tissue Kit (Qiagen, Hilden, Germany) as per the manufacture’s protocol. Briefly, 20 µl of proteinase K was added into the 1.5-ml eppendorf tube with ticks’ homogenates. Buffer ATL was mixed into the mixture and vortexed. Two hundred microliter of absolute ethanol was added and vortexed. The content was transferred into the spin column before being centrifuged at 8000 rpm for 1 minute. The flow-through together with the collection tube was discarded. 500 µl of buffer AW1 was added into the spin-column that was placed into a new collection tube and centrifuged for 1 minute. Again the flow-through together with the collection tube was discarded. 500 µl of buffer AW2 was added into the spin-column that was placed in another new collection tube and centrifuged at 13000 rpm for 3 minutes. Only the flow-through was discarded and the collection tube re-used. Centrifugation of the spin column was done for 3 minutes at 13000 rpm. 200 µl of buffer AE was added into the spin column that was placed into a new 1.5-ml eppendorf tube and incubated at room temperatures for 1 minute. The spin column was centrifuged for 1 minute at 8000 rpm to extract the elute. The concentration and purity of the DNA were evaluated using a Nanodrop spectrophotometer (Thermo Fisher Scientific Wilmington, USA) and stored at −20°C.

#### Molecular detection and identification of tick-borne pathogens

Tick-borne pathogens were screened using HRM and characterized by sequencing of specific RT-PCR products of *Anaplasma, Ehrlichia*, and *Rickettsia* using genus-specific primers (Table 1). For HRM, a 10 µl reaction volume containing final concentration of 1x HOT FIREPol Eva-Green HRM mix (Solis BioDyne, Tartu, Estonia), 500 nM of the respective forward and reverse primers **(Table 1)**, 100 ng of template DNA and 5 μl nuclease free water. DNA extracts from *Anaplasma phagocytophilum, Ehrlichia ruminantium*, and *Ricketia africae* were used as positive controls while nuclease-free water was used as a negative control. Minimum infection rate was calculated using the following formula: [number of pathogen positive tick pools / total number of ticks of that species tested] x 1000. The MIR assumes that only one tick is positive in a pool.

## Results

### Tick diversity and abundance

At near Tsavo National Reserve in Taita Taveta County, a total of 317 (281 adult ticks and 36 nymphs) was collected which includes seven species **(Table 2)**. *Amblyomma* was the most sampled genus with *Amblyomma gemma* being the most sampled species (n=135, 42.6%). Other *Amblyomma* species sampled was *Amblyomma Variegatum* (n= 40, 12.62%). Greatest species diversity was identified in *Rhipicephalus* genus with four species identified that includes; *Rhipicephalus appendiculatus* (n=44, 13.9%), *Rhipicephalus Averts* (n=1, 0.31%), *Rhipicephalus Decoloratus* (n=5, 1.6%), *Rhipicephalus Pulchellus* (n= 91, 28.7%). A single species of *Hyalomma* sp. was sampled **(Table 2)**.

From near Shimba Hill game reserve (SHNR), a total of 240 adult’s ticks were sampled that comprises of eight species **(Table 2)**. *Amblyomma* was the most sampled genus and again *Amblyomma gemma* being the most sampled species (n=156, 65 %). Other *Amblyomma* species sampled includes; *Amblyomma Lepidum* (n= 5, 2.1 %), *Amblyomma Variegatum* (n= 15, 6.3 %). Greatest species diversity was also identified in *Rhipicephalus* genus with four species identified that includes; *Rhipicephalus appendiculatus* (n=18, 7.5 %), *Rhipicephalus Averts* (n=6, 2.5 %), *Rhipicephalus Decoloratus* (n=4, 1.7 %), *Rhipicephalus Pulchellus* (n= 34, 14.2 %). The least sampled species was A single species of *Hyalomma S*cupense (n=2, 0.83 %) as shown in Table 2.

### Tick-borne pathogens identified

At near Tsavo National Reserve (TNR) in Taita Taveta County, a total of three pools of *Rhipicephaline appendiculatus* were positive for *Theileria parva* (GenBank accession Number OL451869-OL451871), two pools of *Rhipicephaline evertsi* for *Anaplasma platys* (GenBank accession Number OL451873-OL451874) and one pool of *Amblyomma variegatum* nymphs for *Rickettsia africae* (GenBank accession Number OL466919) as shown in **Table 3**.

From near Shimba Hill game reserve (SHNR), Kwale County, *Rickettsia africae* pathogen was detected in two pools (GenBank accession Number OL466921-OL466922) of *Am. variegatum* and one pool of *Am. Gemma. Rickettsia sp*. (GenBank accession Number OL466920) and *Anaplasma sp*. (GenBank accession Number OL451872) were detected in *Am. Gemma* and *Rh. evertsi* respective. *R. aeschlimannii* (GenBank accession Number OL466924) was detected in a pool of *Am. Gemma*.

## Discussion

Detection of tick-borne zoonotic diseases at the human-wildlife-livestock interfaces of Shimba Hills National Reserve and the Tsavo National Reserve, gives an insight into the possibility of tick-borne zoonotic diseases spilling over from the wildlife to livestock and humans. Most of the tick’s species identified in this study have been incriminated as the vectors for transmission of tick-borne diseases of public health and veterinary concerns (20).

*Rickettsia africae* is a zoonotic tick-borne pathogen which causes African tick bite fever which manifest with headache, fever, rush, myalgia and skin lesion at the site of tick bite (21). It is often misdiagnosed at the clinical setting because it shares most of the symptoms with other febrile illness like malaria. This calls for robust surveillance and inclusions of *Rickettsia africae* in hospitals and other clinical settings. In this study, *Rickettsia africae* was detected in *Am. gemma* and *Am. variegatum* ticks. These findings are consistence with those that confirms the circulation of *Rickettsia africae* among the *Amblyomma* species (14, 18). Further, *Rickettsia africae* was detected in nymphs of *Am. variegatum* ticks. We can’t tell whether the nymphs got infected through transovarial or direct from the cattle since sampling was done from the cattle.

*Rickettsia aeschlimannii* is also considered to be a zoonotic tick borne pathogen which has been reported on patients travelling from Africa (22). This pathogen has been detected in detected in *Hyalomma* ticks in Kenya (12). It is believed ticks of the genus *Hyalomma* serves as reservoirs for *R. aeschlimannii* (23). However, in the present study, *R. aeschlimannii* was detected in *Am. gemma*. According to our knowledge this is the first time *R. aeschlimannii* has been detected in *Am. gemma*. Migratory birds may play an important role in the epidemiology of *R. aeschlimannii* by carrying *R. aeschlimannii* infetcted ticks from one region to another (24).

Uncharacterized Rickettsia sp. was detected in *Am. variegatum* ticks. Although its pathogenicity was unknown, it can be a public health threat in future (25). Many known *Rickettsia* spps. of public health concern today, were initially considered not to be harmful to humans. This species needs further characterization with better makers such as *ompA, sca4, 17kDa* (14).

*Anaplasma platy* is a TBP infecting dogs which is transmitted by brown dog ticks (*Rhipicephalus sanguineus*) (26). The pathogen has been reported also to infect humans causing mild headache, lethargy and myalgia (27). Humans get infected by coming into contact with dogs infested with ticks infected with *Anaplasms platy* (28). In the current study, *Anaplasma platy* was detected in *Rh. Evertsi*.

*Theileria parva* is a TBP of veterinary importance which causes East Coast Fever in cattle (13). In this current study, *Theileria parva* was detected in *Rh*.*appendiculatus* ticks. These findings were consistent with those of Oundo et al. (14). The cattle get infected by coming into close proximity to buffaloes which are natural reservior for *T*.*parva* (29).

## Conclusion

The detection of *Rickettsia africae, Rickettsia aeschlimannii*, and *Anaplasma platy* gives an insight into the possibility of transmission of zoonotic tick-borne pathogens from wildlife to humans at human-wildlife-livestock interfaces. Since persons infected with these pathogens present with fever like, any febrile illness, it is paramount to include tests that can diagnose these infections at clinical settings.

**Fig.1.**
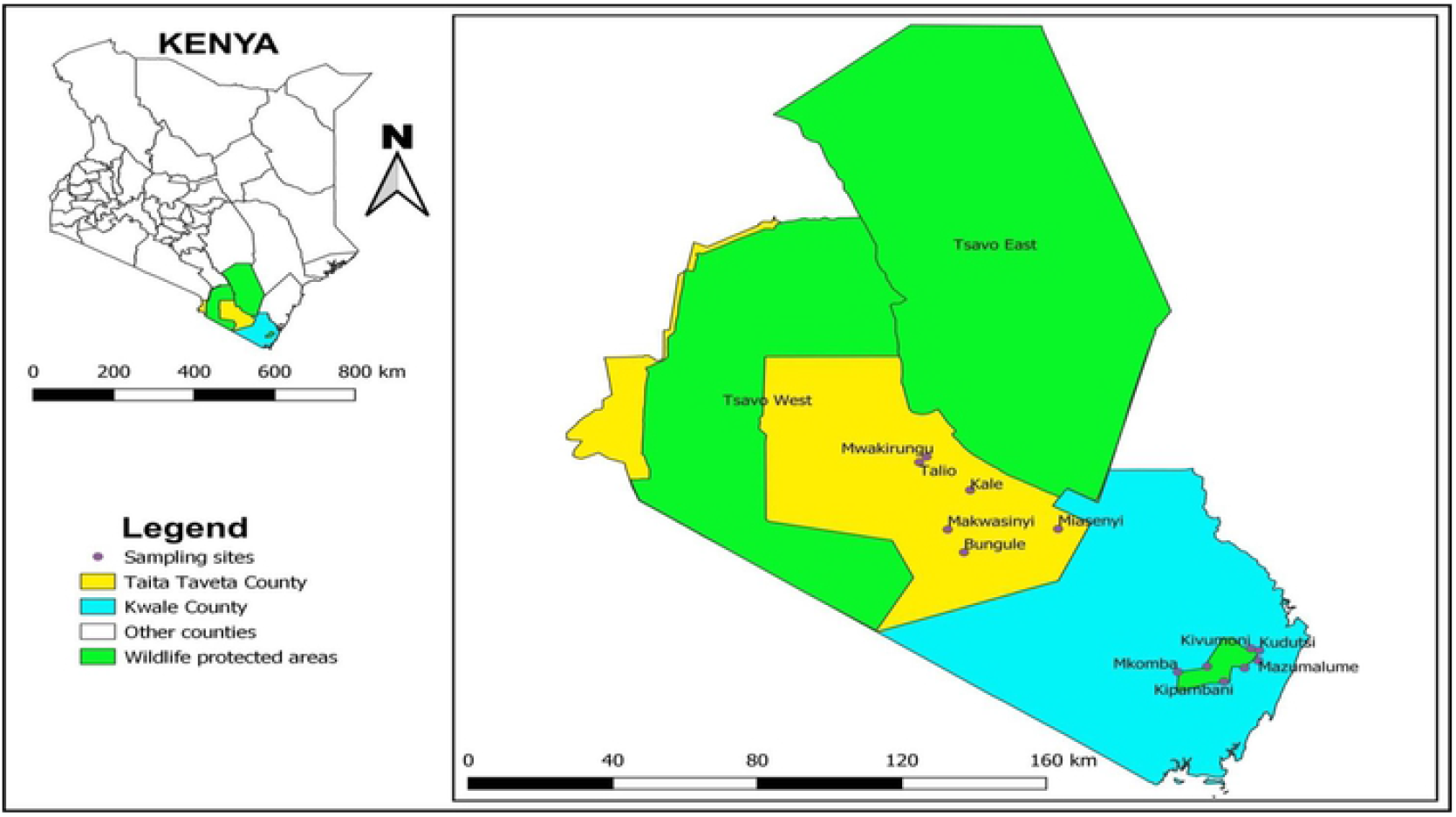
Map of ticks sampling locations near the TNR in Taita Taveta County, Kenya, and near the SHNR in Kwale County, Kenya. The map was prepared using common-license shape files in QGIS software. TNR, Tsavo National Reserve; SHNR, Shimba Hills National Reserve.

## Acknowledgments

I acknowledge Msambweni department of vector borne diseases for their technical assistance.

## References

1. de la Fuente J, Kocan KM. Advances in the identification and characterization of protective antigens for recombinant vaccines against tick infestations. Expert Rev Vaccines. 2003;2: 583–593. doi:10.1586/14760584.2.4.583.

2. Peña A, Jongejan F. Ticks feeding on humans: a review of records on human-biting Ixodoidea with special reference to pathogen transmission. Exp Appl Acarol. 1999;23: 685–715. doi:10.1023/a:1006241108739

3. Herrmann JA, Dahm NM, Ruiz MO, Brown WM. Temporal and Spatial Distribution of Tick-Borne Disease Cases among Humans and Canines in Illinois (2000-2009). Environ Health Insights. 2014;8: 15–27. doi:10.4137/EHI.S16017.

4. Dumitrache MO, Gherman CM, Cozma V, Mircean V, Györke A, Sándor AD, et al. Hard ticks (Ixodidae) in Romania: surveillance, host associations, and possible risks for tick-borne diseases. Parasitol Res. 2012;110: 2067–2070. doi:10.1007/s00436-011-2703-y.

5. Ergonul O. Crimean-Congo hemorrhagic fever virus: new outbreaks, new discoveries. Curr Opin Virol. 2012;2: 215–220. doi:10.1016/j.coviro.2012.03.001.

6. van Nunen S. Tick-induced allergies: mammalian meat allergy, tick anaphylaxis and their significance. Asia Pac Allergy. 2015;5: 3–16. doi:10.5415/apallergy.2015.5.1.3.

7. Gubler DJ. Vector-borne diseases. Rev Sci Tech. 2009;28: 583–588. doi:10.20506/rst.28.2.1904.

8. Zaid T, Ereqat S, Nasereddin A, Al-Jawabreh A, Abdelkader A, Abdeen Z. Molecular characterization of Anaplasma and Ehrlichia in ixodid ticks and reservoir hosts from Palestine: a pilot survey. Vet Med Sci. 2019;5: 230–242. doi:10.1002/vms3.150.

9. Hotez PJ, Kamath A. Neglected tropical diseases in sub-saharan Africa: review of their prevalence, distribution, and disease burden. PLoS Negl Trop Dis. 2009;3: e412. doi:10.1371/journal.pntd.0000412.

10. Leclerque A, Kleespies RG. A Rickettsiella bacterium from the hard tick, Ixodes woodi: molecular taxonomy combining multilocus sequence typing (MLST) with significance testing. PLoS One. 2012;7: e38062. doi:10.1371/journal.pone.0038062.

11. Wesonga FD, Kitala PM, Gathuma JM, Njenga MJ and Ngumi P N. An assessment of tick-borne diseases constraints to livestock production in a smallholder livestock production system in Machakos District, Kenya. Livestock Research for Rural Development. 2010; 22:111. Retrieved January 20, 2022, from http://www.lrrd.org/lrrd22/6/weso22111.htm

12. Omondi D, Masiga DK, Fielding BC, Kariuki E, Ajamma YU, Mwamuye MM, et al. Molecular Detection of Tick-Borne Pathogen Diversities in Ticks from Livestock and Reptiles along the Shores and Adjacent Islands of Lake Victoria and Lake Baringo, Kenya. Front Vet Sci. 2017;4: 73. doi:10.3389/fvets.2017.00073.

13. Walker AR, Bouattour A, Camicas J-L, Preston PM. Ticks of Domestic Animals in Africa: a guide to identification of species. Bioscience Reports, Edinburgh; 2014. Available: http://dx.doi.org/.

14. Oundo JW, Villinger J, Jeneby M, Ong’amo G, Otiende MY, Makhulu EE, et al. Pathogens, endosymbionts, and blood-meal sources of host-seeking ticks in the fast-changing Maasai Mara wildlife ecosystem. PLoS One. 2020;15: e0228366. doi:10.1371/journal.pone.0228366

15. Nijhof AM, Bodaan C, Postigo M, Nieuwenhuijs H, Opsteegh M, Franssen L, et al. Ticks and associated pathogens collected from domestic animals in the Netherlands. Vector Borne Zoonotic Dis. 2007;7: 585–595. doi:10.1089/vbz.2007.0130

16. Roux V, Raoult D. Phylogenetic analysis of members of the genus Rickettsia using the gene encoding the outer-membrane protein rOmpB (ompB). Int J Syst Evol Microbiol. 2000;50 Pt 4: 1449–1455. doi:10.1099/00207713-50-4-1449.

17. Tokarz R, Kapoor V, Samuel JE, Bouyer DH, Briese T, Lipkin WI. Detection of tick-borne pathogens by MassTag polymerase chain reaction. Vector Borne Zoonotic Dis. 2009;9: 147–152. doi:10.1089/vbz.2008.0088..

18. Mwamuye MM, Kariuki E, Omondi D, Kabii J, Odongo D, Masiga D, et al. Novel Rickettsia and emergent tick-borne pathogens: A molecular survey of ticks and tick-borne pathogens in Shimba Hills National Reserve, Kenya. Ticks Tick Borne Dis. 2017;8: 208–218. doi:10.1016/j.ttbdis.2016.09.002.

19. Georges K, Loria GR, Riili S, Greco A, Caracappa S, Jongejan F, et al. Detection of haemoparasites in cattle by reverse line blot hybridisation with a note on the distribution of ticks in Sicily. Vet Parasitol. 2001;99: 273–286. doi:10.1016/s0304-4017(01)00488-5.

20. Parola P, Inokuma H, Camicas JL, Brouqui P, Raoult D. Detection and identification of spotted fever group Rickettsiae and Ehrlichiae in African ticks. Emerg Infect Dis. 2001;7: 1014–1017. doi:10.3201/eid0706.010616.

21. Raoult D, Fournier PE, Fenollar F, Jensenius M, Prioe T, de Pina JJ, et al. Rickettsia africae, a tick-borne pathogen in travelers to sub-Saharan Africa. N Engl J Med. 2001;344: 1504–1510. doi:10.1056/NEJM200105173442003.

22. Parola P, Paddock CD, Socolovschi C, Labruna MB, Mediannikov O, Kernif T, et al. Update on tick-borne rickettsioses around the world: a geographic approach. Clin Microbiol Rev. 2013;26: 657–702. doi:10.1128/CMR.00032-13.

23. Matsumoto K, Parola P, Brouqui P, Raoult D. Rickettsia aeschlimannii in Hyalomma ticks from Corsica. Eur J Clin Microbiol Infect Dis. 2004;23: 732–734. doi:10.1007/s10096-004-1190-9.

24. Rumer L, Graser E, Hillebrand T, Talaska T, Dautel H, Mediannikov O, et al. Rickettsia aeschlimannii in Hyalomma marginatum ticks, Germany. Emerg Infect Dis. 2011;17: 325–326. doi:10.3201/eid1702.100308.

25. Raoult D, Roux V. Rickettsioses as paradigms of new or emerging infectious diseases. Clin Microbiol Rev. 1997;10: 694–719. doi:10.1128/CMR.10.4.694.

26. Abarca K, López J, Perret C, Guerrero J, Godoy P, Veloz A, et al. Anaplasma platys in dogs, Chile. Emerg Infect Dis. 2007;13: 1392–1395. doi:10.3201/eid1309.070021.

27. Arraga-Alvarado CM, Qurollo BA, Parra OC, Berrueta MA, Hegarty BC, Breitschwerdt EB. Case report: Molecular evidence of Anaplasma platys infection in two women from Venezuela. Am J Trop Med Hyg. 2014;91: 1161–1165. doi:10.4269/ajtmh.14-0372.

28. Maggi RG, Mascarelli PE, Havenga LN, Naidoo V, Breitschwerdt EB. Co-infection with Anaplasma platys, Bartonella henselae and Candidatus Mycoplasma haematoparvum in a veterinarian. Parasit Vectors. 2013;6: 103. doi:10.1186/1756-3305-6-103.

29. Magulu E, Kindoro F, Mwega E, Kimera S, Shirima G, Gwakisa P. Detection of carrier state and genetic diversity of Theileria parva in ECF-vaccinated and naturally exposed cattle in Tanzania. Vet Parasitol Reg Stud Reports. 2019;17: 100312. doi:10.1016/j.vprsr.2019.100312

